# Diversification and functional expansion of archaeal TFF machineries

**DOI:** 10.64898/2026.04.22.720090

**Authors:** Shamphavi Sivabalasarma, Sonja-Verena Albers

## Abstract

Archaea have several cell surface structures that belong to the type 4 filament (TFF) superfamily. What all have in common is the presence of core minimal assembly systems consisting of an ATPase, a platform protein, a filament forming protein and a class III signal peptidase. Here we generated novel MacSyFinder2 models to identify and classify archaeal TFF systems. Our analysis revealed a vast diversity of archaeal TFF with several members harboring one or more TFF assembly machineries. Structure-based phylogenetic analyses revealed that the variable N-terminal domain of TFF-related ATPases reflects the subsystem clustering. This indicates a diversification of the core machinery components within the archaeal secretion ATPase family driven through structural innovation. Genome-wide screening of SP-III containing proteins revealed the widespread presence of substrate binding proteins with SPIII. We hypothesize that these binding proteins with canonical SPIII cleavage sites are used to functionalize TFF machineries for efficient substrate scavenging, expanding the functional repertoire of archaeal TFF systems beyond currently characterized roles.

**Author Summary:** The type IV filament superfamily (TFF) comprises a broad group of surface structures that are widespread across both Archaea and Bacteria. While most well-characterized members originate from bacteria, only a limited number of archaeal TFF systems have been experimentally studied so far. Here, we expand the known diversity of archaeal TFF loci through bioinformatic and comparative genomic analyses. Our results show that these systems are far more diverse and versatile than previously appreciated, often exhibiting specialized functions. The structural diversification of the ATPase machinery likely played a key role in driving the functional diversification of TFF systems in Archaea. Overall, these findings deepen our understanding of how archaea adapt and persist in diverse environments, highlighting their surface structures as essential tools for communication, adhesion, and nutrient acquisition.

## Introduction

Archaea harbor versatile cell surface structures that decorate their cell envelope. Many of these structures can be classified within a superfamily of type IV filament structures (TFF)(1). The TFF superfamily comprises various members of cell surface structures in archaea and bacteria such as the type IV pilus machinery or the archaeal motility machinery, the archaellum. This superfamily evolved from a common ancestor with a minimal core machinery: a prepilin/archaellin peptidase, a filament forming subunit and a membrane integral platform protein in interaction with an ATPase (Figure 1). The prepilin peptidase is a class III signal peptidase essential for processing filament subunits that are inserted at the base of a membrane platform protein (2–4). The AAA+ ATPase, in hexameric conformation, provides energy for filament assembly and function(5–9). Each machinery serves a specific function through additional accessory genes at the core gene locus as the archaellum machinery that incorporated stator complexes through duplication of the filament subunit. Archaeal TFF clade members have been studied *in vivo*, revealing their functions as in *Sulfolobus acidocaldarius*, which encodes several TFF machineries. The archaellum machinery, encoded by 7-11 genes, facilitates motility and swimming, while the Aap pilus, is encoded in a 5 gene cluster, supports twitching motility, adhesion, and biofilm formation (6,10). The UV inducible pilus, a TFF specialization hallmark, is induced by UV, enabling cell recognition and DNA exchange for recombination and repair (8,11). Additionally, some methanogens encode the Epd pilus that contributes to cell adhesion (12,13). While all machineries share a similar core machinery, accessory components delineate in each system. The archaellum harbors proteins for the stator complex ensuring torque generation while the UV pilus includes DNA processing proteins in close gene neighborhood (14,15). Previous analyses of putative archaeal TFF machineries clustered these systems in 11 different subsystems differing in their gene locus composition and incorporation of accessory components (16). Here we expanded this analysis on the currently available vast number of archaeal genomes and identified several archaeal organisms encoding at least one archaeal TFF system. MacSyFinder2 (Macromolecular systems finder) was used for a semiautomated search of archaeal TFF machineries. This allowed to discriminate between archaea specific systems and to understand the different composition of machineries of each subsystem (17). Structure-based phylogenetic analysis of the ATPase the core component of the TFF superfamily using FoldTree (18) revealed that they cluster according to the classified subsystem, indicating extensive diversification within the archaeal secretion ATPase family that is observed in the structural variety of the N-terminal domain of the ATPases.

**Figure 1:**
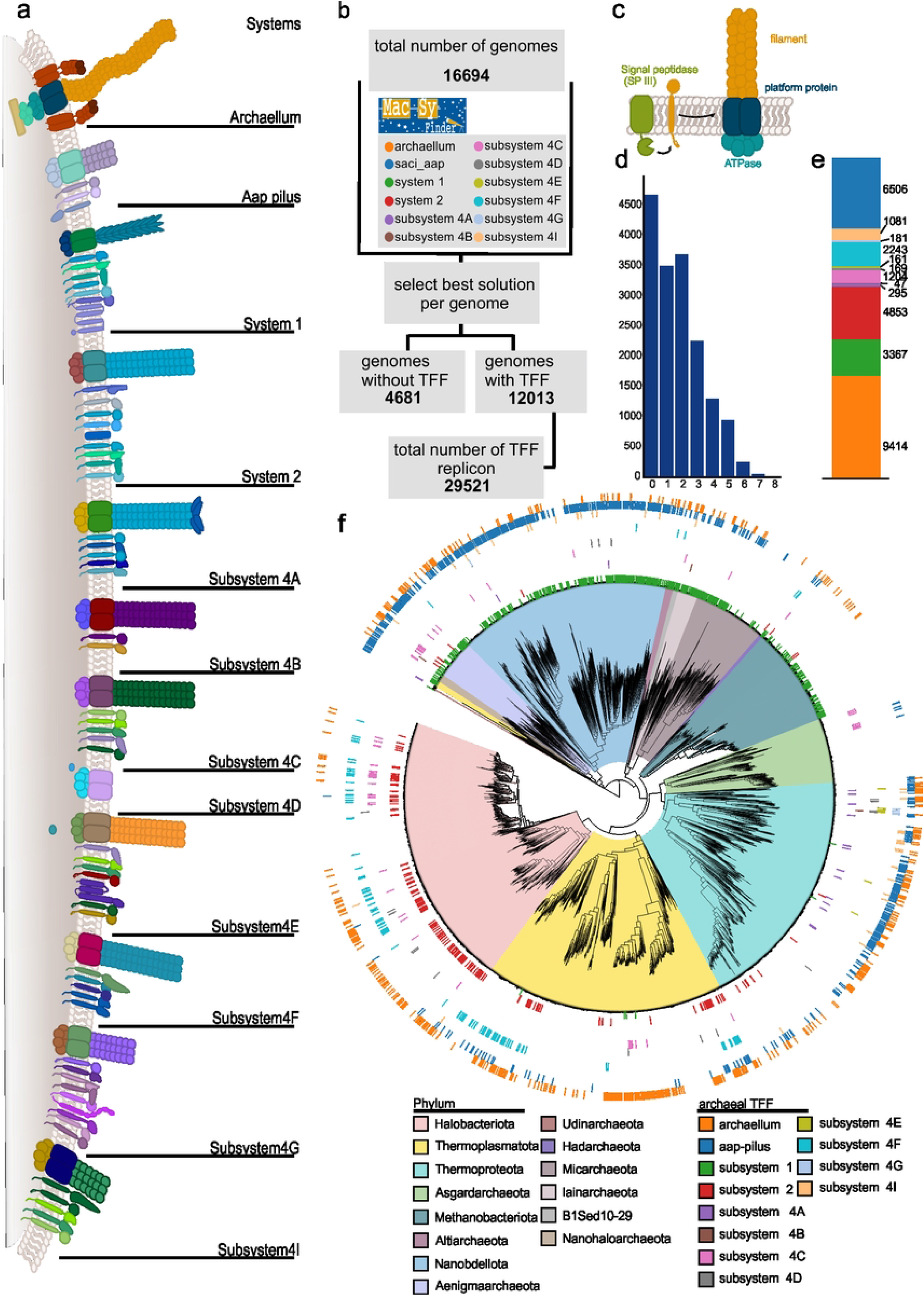
Diversity of archaeal TFF system. **a** schematic displays all 12 archaeal subsystems, each with a macsyfinder model. **b** Workflow of identification of archaeal TFF systems using Macsyfinder2 with models generated for each archaeal TFF. **c** Schematic showing the core machinery components of an archaeal TFF comprising, a class III signal peptidase, a platform protein, an ATPase and a filament forming protein. **d** Number of genomes with encoding multiple TFF loci. **e** Stacked bar graphs showing the number of identified TFF per subsystem. Colors according to **f. f** A GTDB derived phylogenetic tree of archaea with the mapped presence of archaeal TFF subsystems per organisms. Tree scale as indicated.

These novel archaeal surface structures identified might contribute to the efficient functionalization of the cell surface in Archaea, aiding in substrate binding, cell adhesion, cell-cell recognition, and intracellular communication. Indeed, novel substrate-binding proteins were identified through screening archaeal signal peptide sequences for SPIII. These proteins exhibited widespread presence and were processed in a manner similar to TFF subunits, as previously observed for the bindosome in *Saccharolobus solfataricus*(19,20). We propose that certain TFFs serve as the core machinery for incorporating substrate-binding proteins into pilus filaments, ensuring efficient nutrient and substrate uptake expanding the evolutionary and functional landscape of the TFF superfamily.

## Results

### Widespread presence of archaeal TFF clusters in genomes

In a prior study archaeal TFF systems were classified into 11 distinct subsystems by constructing based on sequence similarity of corresponding ATPases (16). Each system was characterized by a specific set of co-localized genes encoding for core machinery and accessory components. Homologs of each component were clustered which in an archaeal cluster of orthologs (arCOG) (Supplementary Table 1, Figure 1a). The arCOGs per systems were extracted and to facilitate detection and differentiation between archaeal TFF systems, models. We generated an internal database of archaeal genomes (16,679 archaeal genomes (as of May 2025)), which was screened for archaeal TFF systems using Macsyfinder2 (17) comprising separate models for archaellum, archaeal adhesion pilus, system 1, system 2 and subsystems 4A-I (Figure1a,b).

We focused on ordered replicons of archaeal TFF systems to differentiate subsystems-based gene co-localization, identifying 29,521 replicons encoding for bona fide TFF machineries (Figure 1b, Supplementary table 1). The ordered replicon search found 12,013 genomes encoding at least one TFF system.

Interestingly, several genomes were found to encode multiple TFF loci in their genomes (Figure 1c), as e.g. the archaellum gene locus that is often accompanied by several other TFF loci in the genome. Although most encode for only one, some seem to harbor up to 8 complete TFF systems in the genome (Figure 1d).

To elucidate the widespread presence of archaeal TFF subsystems, the detected gene loci of ordered replicons per subtype were mapped on the GTDB-derived phylogenetic tree with representative phyla distribution (21) (Figure 1f). The 11 subtypes derived from Makarova et al. (16), consists of the subsystems specific for the archaellum machinery, ups pilus, euryarchaeal specific halo and epd pilus and several other functionally unclassified subsystems. Additionally, a subsystem for the aap pilus was added to distinguish between retractable pili and other nonretractable structures. The majority of classified TFF subsystems are classified as archaella (9414) or archaeal adhesion pilus (6506)(Figure 1e). System 4 consists of a plethora of archaeal TFF machineries with various locus organization and accessory components. According to the presence of accessory components the system 4 is split in 8 subsystems (Subsystem 4A-G and 4I) (Figure 1a)(16). Members of system 4 and its subsystems are found in roughly 2/3 of analyzed replicons and are quite widespread in archaea (Figure 1e). These archaeal TFF systems often occur in multiple loci in genomes as seen in several halophiles or members of Thermococcales however with various sets of co-localized pilins and accessory components in their genomes (Figure 1d, e). Based on phylogenetic analyses archaeal TFF are widespread in most phyla except the Asgardarchaeota (Prometheoarchaeati). Notably a few members of Heimdallarchaea encode a simple archaeal TFF that only contains the core machinery components. *Ca*. Lokiarchaeum ossiferum encodes an archaellum machinery (22) (Figure 1e). Similarly recently characterized Hodarchaeales *Ca*. Margulisarchaeum peptidophila and *Ca*. Flexarchaeum multiprotrusionis as well as *Ca*. Nerearchaeum marumarumayae displayed surface structures that likely belong to the TFF superfamily (23,24). In contrast, the thermoproteal phylum is rich in TFF systems often encoding multiple machineries in their genomes. Euryarchaeota harbor multiple systems as well but typically they belong to system 2, system 1 or subsystem 4C, the latter likely being a halospecific TFF machinery (Figure 1a,e, f). System 1 is predominantly found in methanogens, a few DPANN archaea and thaumarchaeal members (Figure 1f).

#### Analysis of subsystem specific ATPases reveal variable N-terminal domains

All archaeal TFF-related ATPases belong to the family of secretion ATPases comprising members of the bacterial secretion systems and TFF machineries(1,16). The archaeal secretion ATPases group into four arCOGs (arCOG01818, arCOG01817, arCOG01819, and arCOG05609), where ATPases of arCOG05609 are specific to subsystem 4G. We predicted protein structures of ATPases of high-confidence system clusters to reconstruct a structure-based phylogenetic tree using FoldTree (18). The resulting clustering largely mirrors the subsystem organization. Archaellum-related ATPases, as well as those from subsystems 4F, 4G, and 4I, form distinct clusters, reflecting evolutionary diversification of archaeal TFF ATPases and suggesting specialized functions (Figure 2a). Since these ATPases clearly separate from the archaellum ATPase, the corresponding systems are likely unrelated to motility and instead fulfil alternative roles (Figure 2a). In contrast, system 2, Aap-related ATPases, and some subsystems of system 4 cluster together, indicating a similar structural arrangement and possibly related functional mechanisms. This separation of the ATPase tree based on each subsystem points towards on evolutionary diversification of the core machinery components driven through the incorporation of accessory components (Figure 2a). Strikingly, while the core domains remain similar with functional ATPase binding motifs present, additional N-terminal and C-terminal domains can be found (Figure 2b). Their presence is subsystem specific, suggesting that the core machinery components adapted towards their distinct set of accessory proteins and specialized functions within each TFF-related system. We compared the structural predictions of secretion ATPases found in the two model organisms *Sulfolobus acidocaldarius* and *Haloferax volcanii* (Figure 2b). Both encode multiple archaeal TFF systems next to the archaellum-encoding loci (6,8,10,25,26). While the core domains of the ATPase and platform protein remain similar, striking differences can be seen in the N-terminal and C-terminal regions (Figure 2b). The archaellum-related core machinery proteins show the simplest organization while other subsystem related ATPase encode for complex N-terminal or C-terminal domains. These structural differences likely reflect adaptations of the core machinery to subsystem-specific functionalization. We hypothesize that these extra domains serve as potential interaction interfaces to subsystem-specific accessory components. The most minimal ATPases, such as those linked to the archaellum, lack these domains suggesting that the additional loops and helices in other systems could interfere with or replace a potential rotary mechanism characteristic of simpler ATPases (Figure 2b). Indeed, functionally characterized ATPases and TFF machineries in Sulfolobales indicate that different functions and mechanisms correlate with the structural variety observed(5,6,8,10). Similarly, cross-complementation of ATPases and platform proteins is only limited in *Haloferax volcanii*, likely due to mismatching extra domains (27).

**Figure 2:**
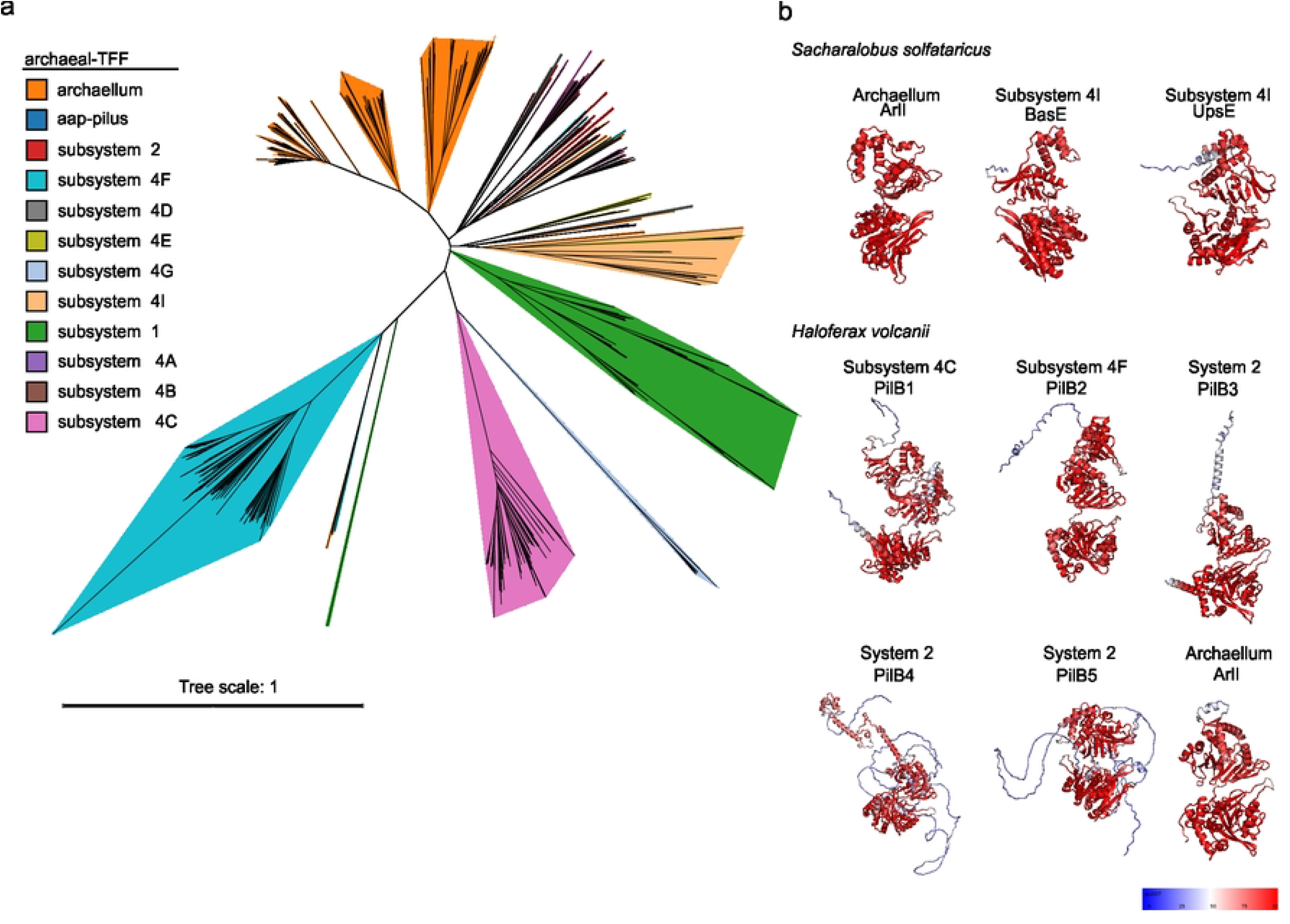
Structural phylogeny of selected TFF related ATPases. **a** Structural phylogenetic tree of ATPases from fold_tree labelled by archaeal TFF system. Tree scale as indicated. **b** Alphafold predicted structures of TFF-related ATPases from the model organism *Sulfolobus acidocaldarius* and *Haloferax volcanii* showing the structural variability across ATPases of different TFF systems. Structures are coloured according to pLDDT value (blue-low, red-high).

#### The archaellum machinery is widespread in archaea

The archaellum, the motility machinery in archaea represents the most widespread TFF identified. 32% of identified replicons were found to encode for the archaellum machinery. The archaellum machinery is often accompanied by additional TFF machineries belonging to other subsystems, suggesting an early specialization of each TFF system towards a dedicated function and limited cross-compatibility between them (Figure 2e). Most archaellum machineries share a conserved core composed of ArlJ and ArlI, representing the platform and ATPase components, respectively (Figure 3). Key accessory proteins unique to and system-defining for the archaellum are the stator proteins ArlF and ArlG. The order of *arlF* and *arlG* together with the presence of either *arlCDE* homologs or *arlX* delineates the archaellum machinery between Thermoproteota and Euryarchaeota (16,28). Most Thermoproteota (except for Thaumarchaeota) encode *arlX*, while euryarchaeal species encode *arlCDE* homologs in different gene fusion or fission arrangements. The presence of *arlCDE* in most genomes coincides with the presence of the archaeal-specific chemotaxis adapter protein CheF, which fits well with experimental findings showing that CheF interacts with ArlCDE(29– 31) (Figure 3). Additionally, homologs belonging to arCOG05057 were identified in some Methanococcales. These encode a S-layer-like protein consisting of several Ig-like folds (Figure 3). This observation coincides with the presence of a similar S-layer-like protein in Thermococcales archaellum loci, indicating that such an association may be more widespread than previously anticipated (16). The resemblance to known archaeal S-layers suggests that this protein could play a role in priming or locally modifying the S-layer to allow the filament to traverse the otherwise rigid and impermeable cell envelope.

**Figure 3:**
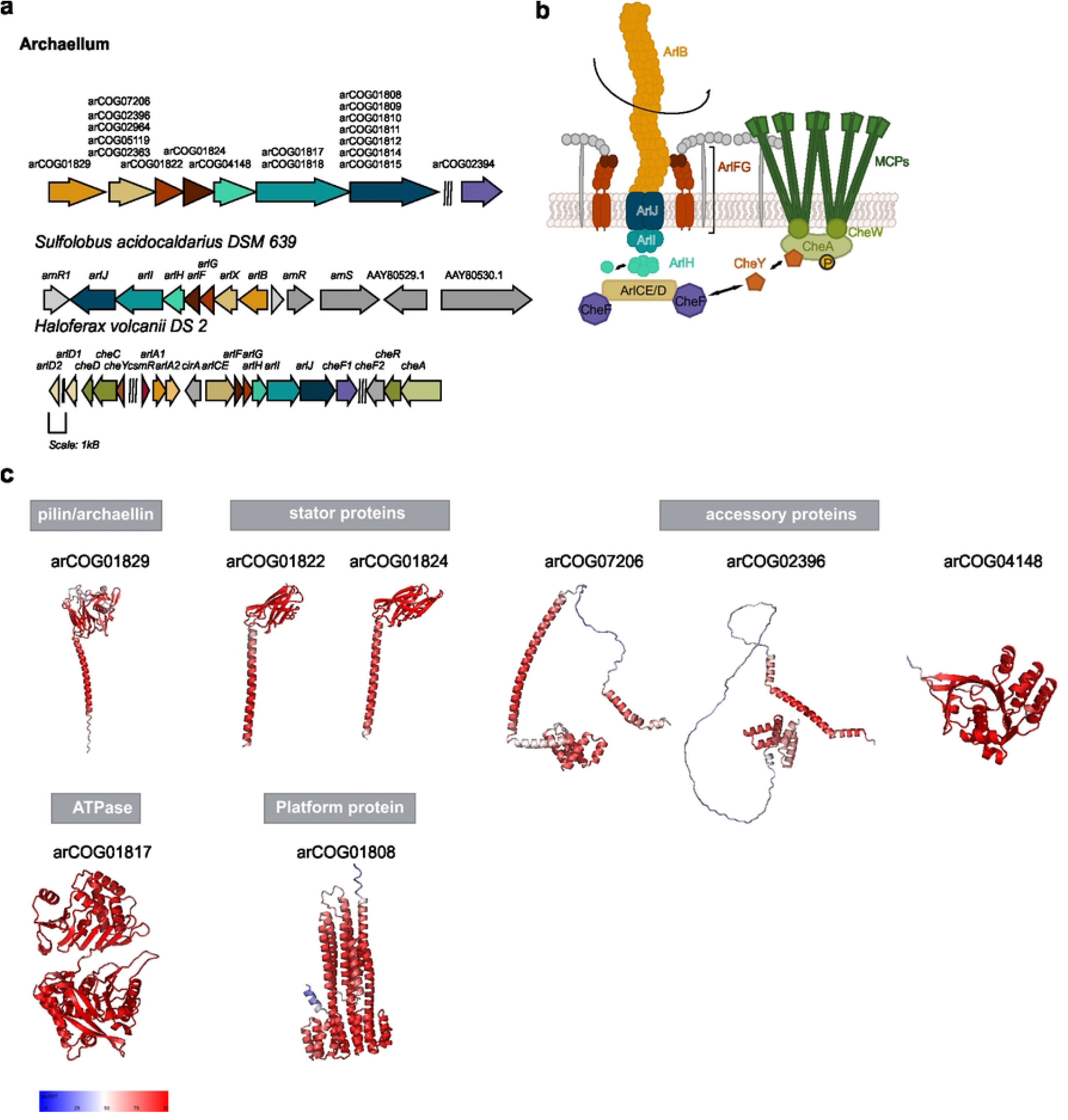
Schematic representation of archaellum TFF. **a** gene locus organization with corresponding arCOGs in exemplary organisms encoding for an archaellum TFF. **b** Schematic representation of the archaellum machinery with its accessory components. **c** Alphafold database derived predicted structures of the core archaellum components. Structures are coloured according to pLDDT value (blue-low, red-high).

#### Aap pilus

The archaeal adhesion pilus is well characterized in Sulfolobales and allows for twitching motility in absence of a dedicated retraction ATPase (7). It mediates surface adhesion and biofilm formation (6) and consists of two pilin homologs, an ATPase, a platform protein, and a protein containing an Fe–S cluster domain (Figure 4). Aap specific pilins belong to arCOG03871 and arCOG03872 (Figure 4a). Structural analyses revealed that a single pilin subunit can adopt three distinct conformations, with varying angles between the hydrophobic N-terminal region and the globular domain, providing filament flexibility necessary for surface-associated motility (32). The Fe–S cluster domain-containing protein (arCOG07320) is found in most *Sulfolobales* encoding an Aap pilus, as well as in some Desulfurococcales *Thermoplasmatales* and *Thaumarchaeota*, including *Nitrososphaera koreensis, Nitrosocaldus uzonensis, Nitrosopumilus maritimus*, and *Ca. Udinarchaeum marinum*, but its function is unknown. In some lineages, such as *Ca*. Jingweiarchaeum tengchongense, multiple Aap loci are present, suggesting a diversification of function. Simpler Aap-like systems lacking the *aapX* homolog are found in several Heimdallarchaeota, Bathyarchaeota, and DPANN members. These minimal systems may still be expressed as *Ignisphaera aggregans*, which encodes a reduced Aap-like machinery, displays surface filaments (33). The Aap-like pilus machineries represent simplified pilus systems that likely function in surface adhesion and possibly in twitching motility, suggesting they may be more widespread across Archaea than previously recognized.

**Figure 4:**
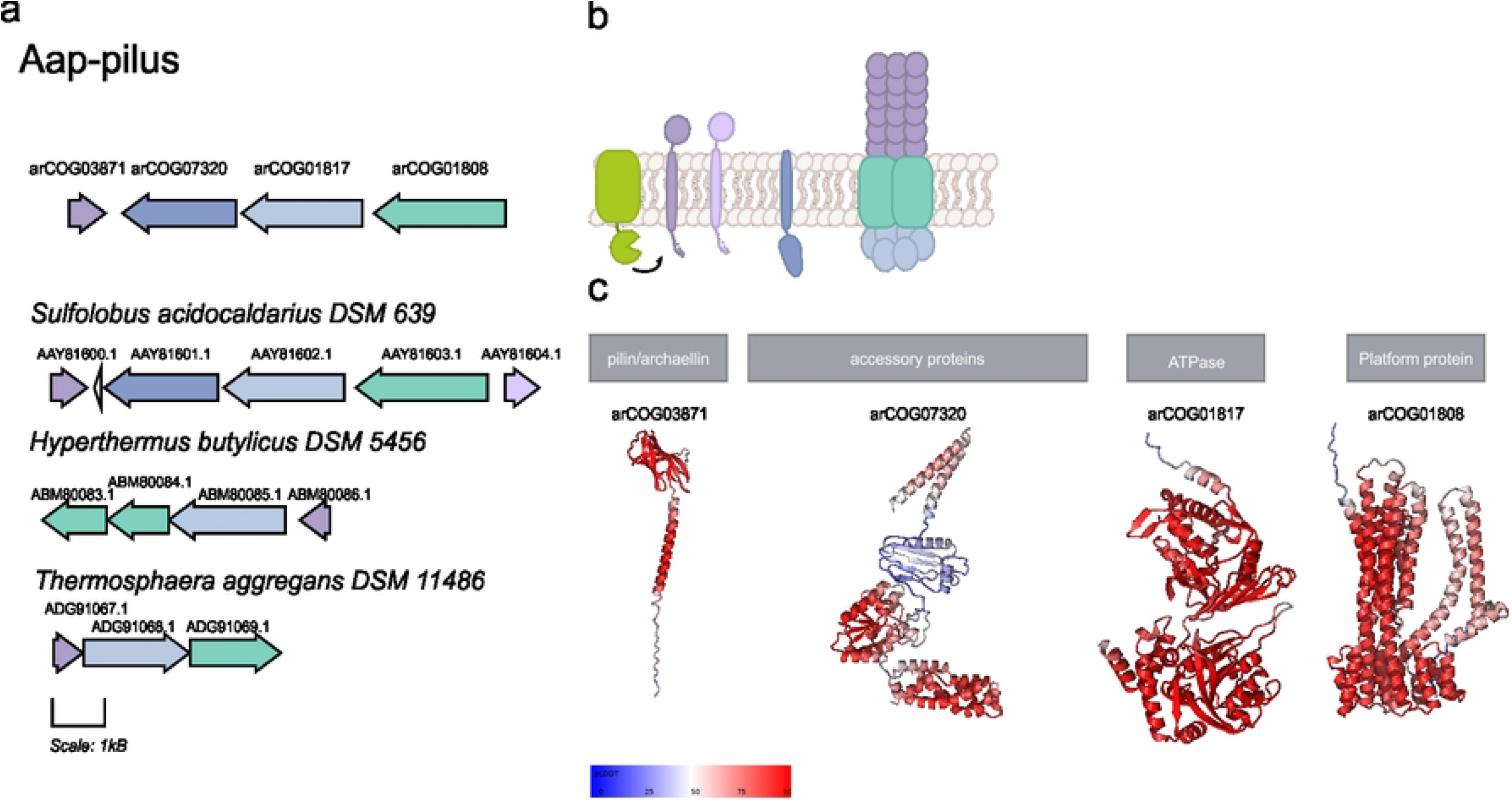
Schematic representation of aap pilus TFF. **a** gene locus organization with corresponding arCOGs in exemplary organisms encoding for an aap TFF. **b** Schematic representation of the aap machinery with its accessory components. **c** Alphafold database derived predicted structures of the aap components. Structures are coloured according to pLDDT value (blue-low, red-high).

#### System 1 is mainly found in methanogens, DPANNs and deep-branching euryarchaeota

Archaeal TFF system 1 clusters are primarily found in Methanococcales, Methanobacteriales, Thermococcales and DPANN archaea. This system includes an ATPase homolog and is characterized by two genes that encode for the platform protein (Supplementary Figure 1). System 1 is characterized by a high variety of pilins and several accessory proteins. The diversity of pilins may support two pili types of system 1: The Epd pilus in Methanococcales, such as *Methanococcus maripaludis* S2 building thin cell surface filaments and an unidentified pilus filament composed of the remaining pilins. Indeed, *Haloferax volcanii* encodes six pilin homologs, all expressed and each one likely forming distinct cell surface filaments (34). The primary Epd-specific pilin from arCOG06620 is missing the C-terminal globular domain, similar to the bacterial Tad pilus, indicating that the bacterial Tad likely originated from the archaeal Epd pilus (1). Additional pilins with a class III signal peptides suggest another system 1 pilus likely expressed under certain conditions. Previously unrecognized system 1 pili in Methanobrevibacter species may aid in adhesion within the gut microbiome or even host gut ephithelia. *Pyrococcus furiosus* and other Thermococcales encode an Epd pilus, highlighting diverse surface structures (Supplementary Figure 1). The Epd pilus contributes to adhesion in *M. maripaludis*. Here, the Epd pilus cluster was extensively studied, and it was shown that the 11 gene cluster is constitutively expressed as one transcriptomic unit. The Epd pilus has its own dedicated signal peptidase, EpdA (arCOG2300)(35). Deleting any of the 11 genes results in nonpiliated cells, while removing minor pilins notably decreases surface pilins (12,36). An adhesin-like domain in mmp0235 and a SPIII protein with a galactose-binding domain suggest roles in surface recognition and adhesion. Standalone pilins at a separate locus are organized with the subsystem 1 pilus. Their deletion in *M. maripaludis* does not affect adhesion, though optimal conditions may be required for the expression of a functional filament (36,37).

#### System 2

Archaeal TFFs belonging to system 2, are predominantly found in Euryarchaeota (Supplementary Figure 2). Often, multiple loci corresponding to subsystem 2 coexist within the same genome; for example, in Halobacteriales, two TFF encoding loci are present, but only the *pil1* locus has been shown to produce functional pili (38). Similarly, in *Haloferax volcanii*, among five *pil* clusters, only *pilBC3* forms visible pili that function in adhesion and biofilm formation (25,27,39,40). The role of the remaining loci is unclear, with similar unidentified loci observed in *Aciduliprofundum boonei* and *Archaeoglobus fulgidus*. These observations suggest that multiple TFF of system 2 loci may coexist in a single genome, potentially serving distinct or condition-specific functions. Makarova et al. hypothesized that system 2 specific TFFs may depend on MinD-like ATPases (16). Indeed, MinD ATPases are widespread and play an essential role in the positioning of the archaellum machinery in H. *volcanii* (41). Furthermore, this suggests a subsystem specific regulation analogous to what is found for the archaellum machinery. System 2 TFFs differ from system 1 by having ATPases with extended N-terminal and sometimes additional C-terminal regions. The platform protein is a single gene, unlike the split in two genes seen in system 1. Multiple pilins are encoded, including SPIII-type and pilin-like proteins without SPIII motifs, alongside membrane proteins likely part of the assembly machinery. In addition, the system 2 loci often encode large adhesin-like proteins, which could be linked to cell adhesion and recognition processes (Supplementary Figure 2). This suggests a potential functional specialization of system 2 pili beyond merely forming filaments. A comparative analysis of organisms with system 2 loci will be required to determine whether additional surface structures co-occur and whether cluster-encoded genes are actively expressed under certain conditions.

#### TFF clusters belonging to System 4

The clusters assigned to subsystem 4 are highly diverse and can be subdivided into several subsystems (4A–4I) that differ substantially in gene neighborhood organization around the homolog of the platform protein and the ATPase (16). The order and domain architecture show striking variability across subsystems, suggesting frequent gene rearrangements and incorporation of accessory domains during evolution. Below, each subsystem is described in more detail:

#### Subsystem 4A and 4B

Subsystem 4A is predominantly found in Thermoproteota, some Nanoarchaeota and Bathyarchaeota, as well as in a few euryarchaeal genomes. The cluster organization is distinct where genes encoding for accessory components are found downstream of the gene encoding for the platform protein (Supplementary Figure 3 a-c). In some DPANN archaea, this includes two platform proteins, as system 1 loci. The minimal subsystem 4A has two platform proteins, an ATPase, a pilin, and often an adhesin (Supplementary Figure 3). 4A specific pilins belong to arCOG11238 and arCOG06117 and a subsystem 4AB specific adhesin that is remotely homologous to the bacterial PilY. In bacteria, PilY is suggested to play a role in adhesion and mechanosensing, linking type IV pili dynamics to intracellular regulation through cyclic nucleotide levels (42,43). In *Pseudomonas aeruginosa*, PilY forms a corkscrew-shaped structure at the secretin, gating the filament assembly and regulating the dynamic cycles of extension and retraction (42).

Subsystem 4B has a relatively simple organization comprising only two pilins encoded with a platform protein and ATPase encoding genes. However, this subsystem comprises two genes encoding for a platform protein as seen for system 1 and subsystem 4A TFFs flanking the ATPase gene up- and downstream (Supplementary Figure 3d-f). This subclade contains arCOG07037 near the major pilin, which encodes an adhesin-like protein that strongly resembles bacterial hook proteins consisting of several Ig domain folds suggesting a role in cell attachment (Supplementary Figure 3f). Subsystem 4B TFFs are restricted to a few lineages, including *Sulfolobus islandicus, S. caldissimus, S. neutrophilus*, and *Thermoproteus uzoniensis*. In *Thermoproteus tenax*, the 4B clade clusters are associated with RecA-like ATPase, CBS-domain proteins, and a eukaryotic DEATH domain protein, which seems to replace KaiB found in some subsystem 4B loci (16). This suggests functional coupling between the TFF assembly and cellular regulatory networks as seen for the complex regulatory network of the archaellum machinery (reviewed in(44)).

#### Subsystems 4C and 4D

The gene loci belonging to the 4C subsystem are found predominantly in halophilic archaea often co-occurring with subsystems of 4F and 4G. Each system encodes a platform protein and an ATPase decorated with accessory pilins and membrane inserted proteins. Distinctive extra loops are found at the N and C-termini on the platform protein and ATPase that are domain specific and might play a crucial role in the function of the archaeal T4P clades. Associated pilins have a variability in their head domain that could indicate special functionalization such as cell adhesion or cell recognition (Supplementary Figure 4 a-c). Similarly, as to subsystem 2, this subsystem seems specific to halophilic euryarchaea yet classifies as its own subsystem due to the gene arrangement and significantly different pilins and accessory proteins. Makarova et al. propose a hybrid origin for this pilin system, with membrane machinery resembling thermoproteal members and pilins likely of euryarchaeal origin(16). This suggests that incorporating base machinery from different members can create functional systems if core components are compatible. In contrast to the previously mentioned clades, Subsystem 4D is notably distinct as it comprises an ATPase, a platform protein but no putative pilin homolog. The presence of a TFIIB-like transcription factor specific for Pyrobaculum and Thermoproteus, as well as an RNA polymerase nearby, implies that it may be a remnant of the TFF system with a repurposed function rather than being involved in pilus assembly (Supplementary Figure 5d-f)(16).

#### Subsystems 4E and 4F

Members of Thermoproteota specifically belonging to Pyrobaculum and Vulcanisaeta encode for a subsystem 4E specific pilus machinery. Its core machinery is likely linked to arCOG03739 and arCOG03740 pilins near the TFF gene locus (Supplementary Figure 6a-c)(16). The gene locus is split into membrane-embedded machinery genes and distantly located pilin-encoding genes (Supplementary Figure 6a-c). Functional expression indicating an assembled filament of corresponding system was shown for *P. arsenaticum* 2GA which forms long thin filaments that serve as attachments for archaeal viruses (45). The pilin, which has an extra C-terminal domain and cysteines forming bonds, likely enhances stability in hyperthermophilic environments. Subsystem 4F resembles subsystem 4C in gene arrangement and distribution, is mainly found in Halobacteria that share the hybrid nature in gene arrangement. These clusters, similar to those of subsystem 4C, have hybrid features with the core machinery, such as the thermoproteal version, while accessory proteins and pilins likely come from euryarchaeal member (16). The 4F gene locus co-localizes either with the archaeal tubulin homolog CetZ2, a CBS domain protein, or an SMC homolog (Supplementary Figure 6d-f). This subsystem contains an adhesin-like protein homolog as well (arCOG02945). These genes found in the gene neighborhood might play a crucial role in the positioning and regulation of the subsystem 4F clusters.

#### Subsystems 4G and 4I

Subsystem 4G is highly complex as it encodes for several pilins, membrane inserted accessory proteins and all systems co-occur with a MinD-like ATPase (Supplementary Figure 7). These subsystems are mainly found in Pyrobaculum and Thermoproteus, often co-occurring with a subsystem 4E TFF locus. The hallmark of this cluster is, next to the several TFF associated components, the presence of large adhesin-like proteins that are rich in Ig folds (arCOG05616). These structured Ig folds resemble the prokaryal hook like proteins recently characterized in *Aureispira* sp., also found in some members of the Chloroflexota phylum (46,47). In *Aureispira* sp., these hook-like proteins facilitate interaction with prey flagella, enabling ixotrophic behavior. Their function in archaea remains unknown. Similar to their role in bacteria, these proteins might be involved in adhesion and/or cell-cell recognition. In gut methanogens, adhesins are crucial for the interaction between secreted membrane vesicles and the host gut epithel cells (48).

Subsystem 4I exclusively includes the bindosome and ups pili of Sulfolobales. Both systems were experimentally studied and shown to be expressed in members of the Sulfolobales (8,20,49–51). The Ups pilus is encoded in a gene locus containing two pilins, an unknown protein consisting of several Ig folds (UpsX, arCOG05986) and its corresponding membrane platform protein and ATPase (Supplementary Figure 7ab). The Ups pilus is induced by UV stress and mediates species specific cell-cell interaction for Ced system mediated DNA exchange and DNA repair using homologous recombination (8,11). Owing to its function, the gene neighborhood of the ups pilus is highly conserved and contains a helicase, endonuclease, and a glycosyltransferase that might play a role in DNA exchange (15). In contrast to the ups pilus, the bindosome is characterized by three pilins that are found in close proximity to the membrane spanning assembly machinery encoded by *basEF* (51) (Supplementary Figure 7de).

Several Sulfolobales members use sugar scavenging bindosome machinery. These bindosomes are associated with an ABC transporter and substrate binding protein whose N-terminus follows the TFF pilin blueprint (20,51,52). While the core machinery is found in one gene locus, the sugar binding proteins are distantly located in the genome, mostly as part of the ABC transporter locus. The sugar binding proteins that are processed by the class III peptidase ensure efficient nutrient uptake through low-affinity sugar binding (52). This shows that base machinery and binding proteins are regulated differently and adapt to substrate availability, with the sugar binding proteins induced by the presence of specific sugars (53). Deletion of the core machinery of the bindosome substantially reduces glucose uptake through its specific binding protein. This indicates that TFF base machinery, namely BasEF in Sulfolobales serves as the base machinery for the sugar binding proteins such as GlcS, AraS, and TreS, however, growth is not delayed (19,20,53).

### Widespread presence of bindosomes with a broad substrate range in archaea

Class III signal peptide proteins include both TFF-related pilin and substrate binding proteins, mainly for sugars, in Sulfolobales like *Saccharolobus solfataricus*, which has about seven sugar-binding proteins (35). These proteins share a conserved N-terminus but diverse C-terminal regions due to varying binding properties. They often co-localize with ABC transporters rather than TFF clusters, indicating regulation based on substrate availability. Despite having 8 binding proteins, *S. solfataricus* has only one substrate binding related TFF machinery, likely serving multiple binding proteins depending on available substrates. To elucidate whether more archaea functionalize a TFF for substrate binding, archaeal genomes were screened for proteins with a class III signal peptide. Candidate proteins were analyzed using hhmscan that later allowed selection of proteins with potential substrate binding function (Figure 5). Possible signal peptides with a lipobox motif indicating a cleavage with signal peptidase II and TAT substrates were excluded, resulting in 4364 potential binding proteins with a class III signal peptide (Figure 5a). Interestingly, these proteins do not only reside with Sulfolobales but are also found in several euryarchaeal members. Analysis of the frequency of binding proteins per genome showed that most genomes contained one or two binding proteins with a SPIII cleavage site, but genomes with up to 9 binding proteins were also found (Figure 5c). Around 3002 binding proteins were found in genomes that also contained a TFF machinery that could serve as the base for the substrate binding proteins. Although approximately 1355 genomes that contained a class III signal peptide with a binding protein did not encode a TFF locus, 1019 genomes did contain a putative SPIII peptidase. This peptidase would enable the correct processing of the substrate binding protein. (Figure 5a).

**Figure 5:**
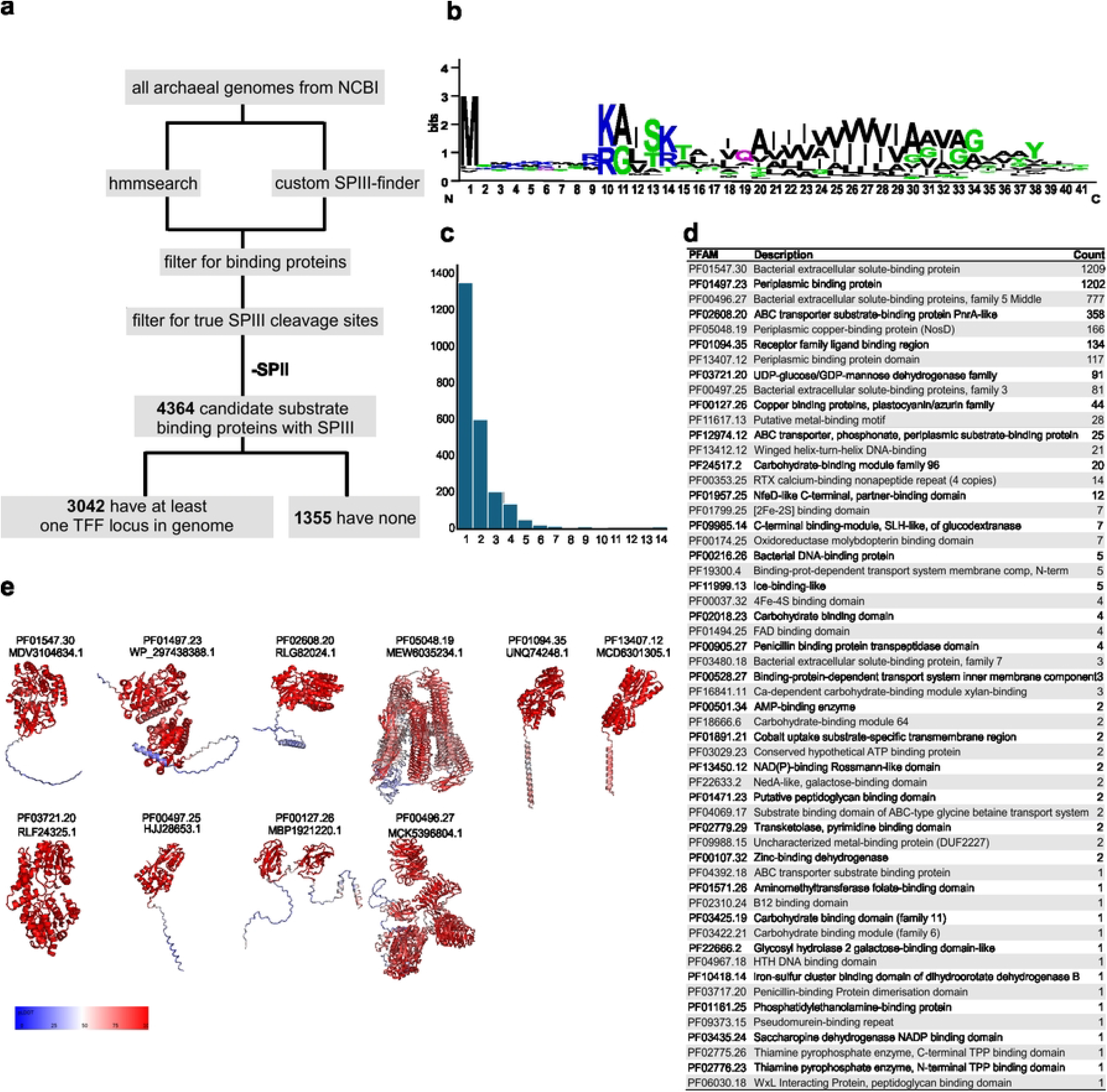
Identification of putative binding proteins with a SPIII. **a** Schematic representation of the identification of binding proteins with SPIII. **b** Weblogo of N-termini of identified binding proteins with conserved SPIII cleavage site and hydrophobic domain. **c** Frequency of identified binding proteins per genome. **d** All PFAM domain identified from hmmsearch of binding proteins. **e** Alphafold3 predicted structure of top 10 PFAM domains showing variable c-terminal domain organization and a hydrophobic N-terminal helix. Structures are coloured according to pLDDT value (blue-low, red-high).

We mapped the PFAM domains identified through hmmscan to putative binding proteins using SPIII. The top ten candidates suggest that nearly all of them are involved in substrate binding, which is commonly associated with ABC transporters. (Figure 5d). For most, the exact substrate is unknown, however, it can be assumed that the binding capability ranges from sugars to peptides and even specific molecules and metal ions. The prediction of the structure of exemplary binding proteins per PFAM shows a variety of C-terminal domain architectures ranging from small length globular folds to long beta-helical structures (Figure 5e). All of them have a hydrophobic TM-domain in common, which precedes the functional domain of the binding protein. The presence of this TM domain, along with the SPIII cleavage site, in a manner similar to pilins and archaellins is striking. This suggests that it is likely inserted into a TFF-pilus structure, which would facilitate substrate scavenging in archaea. Alignment of all extracted N-termini shows highly conserved positive charges preceding the hydrophobic domain that are consistent with the well-studied class III signal peptide (Figure 5b).

## Discussion

### Widespread diversity of archaeal TFF machineries

Screening of archaeal type IV filament (TFF) systems shows a remarkable diversity, identifying 29,521 TFF-related loci and several genomes with multiple systems. A common feature among all systems is the presence of the essential components defining a TFF machinery: a single ATPase, a membrane platform protein, a class III signal peptidase, and at least one pilin responsible for filament formation. Each subsystem incorporates additional accessory proteins that are unique and critical to its respective functions. While multiple pilin genes and platform protein genes may exist within a single locus, the ATPase gene remains in a single copy per subsystem. There is no evidence of gene fission or fusion events for ATPases, contrasting with bacterial TFF systems where ATPase duplication has occurred for different functions (1). In archaea, however, there is considerable variability among pilins, both major and minor, some of which are located within subsystem loci and others as standalone genes. This suggests that the current model used by MacSyFinder2 may not fully capture the total number of potential pilins through the ordered search approach. Indeed, while ordered replicons encoding a TFF locus-like operon organization only yielded the detection of core machinery components in most asgardarchaeal genomes, an unordered search revealed the presence of potential pilins in these genomes. It is tempting to speculate whether pilins and core machinery components were acquired independently of each other.

Genomes frequently contain more than one TFF locus, and both pilin and the core machinery gene structure indicate specialization and functionalization rather than redundancy. For example, in *Haloferax volcanii*, cross-complementation experiments demonstrated that the base machinery from one TFF locus cannot rescue deletion phenotypes in another, indicating functional divergence (27). Comparative analysis of ATPase and platform proteins in genomes with multiple systems, such as *H. volcanii* and *S. solfataricus*, reveals significant structural divergence, particularly in the N- and C-terminal domains of ATPases, which further limits cross-complementation (Figure 2). Structure-guided phylogenetic analysis demonstrates that ATPases group according to their associated TFF systems, reflecting structural diversification driven by adaptation to distinct cellular functions. Many subsystems include proteins with multiple Ig-folds, adhesin-like domains, or beta-helical sheets. For instance, subsystem 2 contains a PilY-like adhesin implicated in cell adhesion and surface recognition, while the functions of similar domains in other subsystems remain unclear. The structural resemblance of some to archaeal S-layer proteins suggests a role in priming the cell wall allowing the pilus filament to traverse the envelope. Notably, modeling of S-layer and surface filaments of *S. acidocaldarius* indicates that local S-layer rearrangements are required to facilitate pilus insertion (54).

### Functionalized TFF Machineries for substrate binding and nutrient scavenging

Substrate-binding proteins with class III signal peptides (SPIII) are widely distributed, initially identified in Sulfolobales but found in other archaeal lineages as well(35) (Figure 5). These proteins are characterized by highly conserved N-termini with class III signal peptides and a hydrophobic N-terminal alpha helix, which might facilitate integration into a possible pilus structure. Alpha-helical interactions stabilize the protein within the filament and are critical to filament architecture. In bacterial systems, pilus filaments are rarely composed of a single pilin; minor pilins and tip complexes contribute to functional diversification such as DNA uptake and adhesion for cell flocculation (55–57). These observations support the hypothesis that a base TFF machinery may serve as a versatile platform for multiple substrate-binding proteins for efficient substrate binding and uptake via the corresponding ABC transporter. In *S. solfataricus*, sugar-binding proteins are induced upon presence of corresponding sugar and their sugar-scavenging activity relies on the bindosome base structure (19,20,53). Indeed, surface proteome studies in *Aeropyrum pernix* showed the expression of a substrate binding protein with a class III signal peptidase indicating its presence and assembly outside of the Sulfolobales (58). The ability of TFF subsystems to accommodate various substrate-binding proteins suggests an expanded functional range of TFF members beyond motility, adhesion and sugar uptake. This diversity is seen in the structural variability of the N-terminal ATPase domain likely arose through extensive evolutionary diversification and incorporation of novel components, which have been repurposed for new functions (Figure 5). While the core machinery proteins (ATPase and platform protein) retain structural similarity, additional domains have evolved in each subsystem, limiting cross-complementation and hinting at specialization for distinct functions. Archaea occupy a wide variety of environments and ecological niches, and the variability in TFF systems reflects this diversity. Future studies integrating genomics, structural biology, and *in vivo* localization will be essential to fully elucidate the novel functions of TFF machineries beyond those currently characterized.

## Materials and Methods

### Identification of TFF machineries

Archaeal Type IV filament systems were identified in archaeal genomes using MacSyFinder2(17). A total of 16,694 archaeal genomes (NCBI, release May 2025) were screened using a set of custom MacSyFinder models specifically designed to capture the diversity of archaeal TFF subsystems(17). MacSymodels per archaeal TFF subsystem according to Makarova et al., 2016 were generated(16). An additional model for the archaea specific adhesion pilus was incorporated to specifically classify it as a separate system. Each subsystem model included the core components: ATPase, platform protein, prepilin peptidase as mandatory components, and the remaining pilins and accessory proteins were classified as accessory. MacSyFinder was run in the “ordered replicon” mode using a custom wrapper script to facilitate detection of co-localized genes within a genomic locus for robust subsystem differentiation. Asgardarchaeal genomes, as well as Thermoplasmata and Nanobdellati were specifically screened using the unordered replicon mode to analyze the presence of candidate genes related to TFF.

### Analysis of gene locus

The exemplary gene locus per system from MacSyFinder2 outputs was visualized with Gene Graphics (Harrison et al., 2018).

### Structure guided phylogenetic tree

ATPases of high confidence TFF clusters were extracted and uniprot identifiers and corresponding Alphafold predicted structures were retrieved using a custom python script that was written with help of the GPT-4.1 AI model. Fold_tree 1.0.0(18) was used to construct a structure guided phylogenetic tree using the command *snakemake --cores 24 --use-conda -s ./workflow/fold_tree --config folder=./myfam filter=False custom_structs=True*.

### Mapping archaeal TFFs onto a reference phylogenetic tree

A reference phylogenetic tree of archaeal genomes was derived from GTDB (release 10) (May 2025)(21). The detected TFF systems were mapped onto this phylogenetic tree to visualize their distribution in archaeal lineages. The mapping was performed using iTOL (Interactive Tree of Life) with colour-coded annotations for each subsystem and the number of TFF loci per genome (59).

### Identification of binding proteins with SPIII

SPIII-containing proteins were identified using a custom script identifying archaeal signal peptides, that was written with help of the GPT-4.1 AI model. This script combines screening for N-terminal transmembrane helices with TMHMM 2.0 with SPIII specific cleavage motif according to Szabo et al., 2007 (35). Additionally, potential N-termini containing a Tat secretion sequence or a lipobox were screened. Proteins containing a TM segment within the first 30 amino acids and a SPIII cleavage site were retained for further analysis. Subsequently, HMMERv3.4 (hmmsearch) was used to identify conserved domains associated with SPIII-bearing proteins using PFAM database(60). The predicted structures were retrieved from the Alphafold database and modelled using PyMol.

### AI tool statement

During the preparation of this chapter ChatGPT was used in order to correct the English language. After using this tool/service, the chapter was reviewed and the content was edited as needed taking full responsibility for the content of the publication.

## Acknowledgements

We thank Ute Hoffmann for critical reading of the manuscript and fruitful discussions. S. Sivabalasarma and S.-V.A. were supported by the Collaborative Research Centre SFB1381 funded by DFG (German Research Foundation)—project ID 403222702—SFB 1381. We further acknowledge support from the German Research Foundation (Germany’s Excellence Strategy CIBSS-EXC-2189, project ID 390939984 to S.V.A.).

## Data availability

All used scripts and general workflow are available in the github repository (https://github.com/shamphavis/Identification-of-archaeal-TFF-systems-using-macsysfinder2). Data that support the findings of this study are available at doi:10.5281/zenodo.19336109.

